# Clade III HIPP genes encode plasmodesmata-targeted proteins with pleiotropic functions in regulating plant development

**DOI:** 10.64898/2026.06.08.730823

**Authors:** Georgeta Leonte, Cristina Belén Aucapiña, Henriette Weber, Isabel Bartrina, Ondřej Novák, Tomáš Werner, Alicja Marta Górska

**Author notes:** **Corresponding authors:** Tomáš Werner, Alicja M. Górska.

## Abstract

Heavy metal-associated isoprenylated plant proteins (HIPPs) are encoded by large gene families, which have diversified specifically in vascular plants. Their physiological functions and molecular mode of activity are currently largely unknown. In this study, we characterize a group of phylogenetically closely related genes *HIPP32*, *HIPP33,* and *HIPP34* in *Arabidopsis thaliana*, revealing their essential roles in controlling diverse developmental pathways. Through comprehensive genetic analyses, we demonstrate that these genes exhibit partially overlapping pleiotropic functions, influencing multiple aspects of plant growth such as embryogenesis, maintenance of apical meristems, root architecture, shoot branching, leaf morphogenesis and floral organ formation. Transcriptomic profiling of *hipp* mutants identified significant deregulation in several regulatory pathways involved in plant hormone responses, with a specific impact on auxin signaling processes. Interestingly, we show that the analyzed HIPP proteins localize very specifically to plasmodesmata, suggesting their potential function in regulating intercellular communication in shaping plant development.

## INTRODUCTION

Heavy metal-associated isoprenylated plant proteins (HIPPs) constitute one of two plant-specific metallochaperone-like protein families characterized by the presence of a heavy metal-associated (HMA) domain (Dykema *et al*., 1999, Tehseen *et al*., 2010, de Abreu-Neto *et al*., 2013). The HMA domain contains two conserved cysteine residues arranged in a CXXC motif (where ‘X’ denotes any amino acid), which are critical for coordinating heavy metal ions (Suzuki *et al*., 2002, Zschiesche *et al*., 2015). Structurally, the HMA domain is a highly conserved module of approximately 70 amino acids found across microbes, plants, and animals. It adopts a ferredoxin-like βαββαβ sandwich fold, forming a compact metal-binding structure (Eriksson & Sahlman, 1993). HIPPs, along with a second group of metallochaperones known as heavy metal-associated plant proteins (HPPs), are defined by the presence of one or two HMA domains. In addition to metal binding, the HMA domain has been shown to mediate protein–protein interactions (Barth *et al*., 2009), highlighting its functional versatility. A key feature distinguishing HIPPs from HPPs is a C-terminal isoprenylation site, a CaaX motif (where ‘a’ represent an aliphatic residues and ‘X’ can be any amino acid). The CaaX motif is recognized by protein prenyl transferases, which attach a hydrophobic isoprenyl farnesyl (C15) or geranylgeranyl (C20) group to the cysteine residue via a thioether bond. Following prenylation, the -aaX tripeptide is cleaved, and the exposed cysteine is carboxymethylated, resulting in an isoprenylated, methylated C-terminus (Hála & Žárský, 2019). Protein prenylation has been shown to enhance membrane association either through direct intercalation of the prenyl group into the lipid bilayer or via interaction with membrane-associated proteins (reviewed by Tian & Wang, 2025). Importantly, the isoprenylation has been shown to be essential for the plasmodesmata (PD) localization of HIPP proteins (Cowan *et al*., 2018, Guo *et al*., 2021, Were *et al*., 2025). Although the HMA domain and isoprenylation motif are found in proteins across many organisms, their occurrence within the same protein is unique to vascular plants.

Recently, Johnston et al., (2023) identified HIPP proteins among conserved orthogroups represented in PD-associated proteomes. To date, up to 11 HIPP family members have been detected in plasmodesmal proteomic datasets (reviewed by Barr *et al*., 2023). The PD localization of HIPP1 and HIPP7 from *Arabidopsis thaliana*, as well as HIPP26 from *Nicotiana benthamiana*, has been confirmed using live cell imaging (Cowan et al., 2018, Guo et al., 2021). Interestingly, HPP proteins have also been identified in PD proteomes (reviewed by Barr et al., 2023), suggesting the existence of additional PD-targeting signals for HMA domain-containing proteins beyond isoprenylation. In addition to their localization at PD, many HIPP proteins show dual or diverse localization patterns, including the cytoplasm, nucleus, and plasma membrane (Barth et al., 2009, de Abreu-Neto et al., 2013, Cowan et al., 2018, Guo et al., 2021, Zhang *et al*., 2020), suggesting functional diversity within the HIPP family.

To date, most studies on HIPP proteins have focused on their roles in plant responses to abiotic and biotic stresses. In particular, HIPP proteins are frequently discussed in the context of plant-pathogen interactions and have been shown to be direct targets of pathogen-derived molecules, including secreted effector proteins and viral movement proteins (reviewed by Turley & Faulkner, 2025). For example, Cowan et al. (2018) showed that *Nicotiana benthamiana* HIPP26 interacts with the *Potato mop-top virus* (PMTV) movement protein TGB1, which is essential for the viral long-distance movement. NbHIPP26 was shown to localize to the nucleus, plasma membrane, and PD; however, when co-expressed with TGB1, it predominantly localizes to nucleoli and microtubules. The HIPP26-TGB1 interaction is proposed to facilitate long-distance viral movement and enhance drought tolerance via the activation of specific transcriptional responses (Cowan et al., 2018).

In addition to their roles in plant-pathogen interactions, an increasing body of evidence supports the involvement of HIPP proteins in heavy metal tolerance. Numerous studies have demonstrated the metal-binding capacity of HIPPs, primarily through *in vitro* assays using heterologously expressed HIPP proteins. For instance, HIPP7 binds Cu²⁺, Ni²⁺, and Zn²⁺ (Dykema et al., 1999); HIPP6 binds Cd²⁺, Hg²⁺, and Cu²⁺ (Suzuki et al., 2002); HIPP26 binds Cd²⁺, Cu²⁺, and Pb²⁺ (Gao *et al*., 2009); while HIPP3 exhibits significant binding only to Zn²⁺ (Zschiesche et al., 2015). The connection between HIPPs and heavy metal regulation is further highlighted by HIPP transcriptional responses to various metal ions. For instance, expression of Arabidopsis *HIPP6* is upregulated by Hg, Fe and Cu (Suzuki et al., 2002), while Cu and Cd induce *HIPP16* expression in pak choi (Niu *et al*., 2021). Additional *HIPP* genes responsive to heavy metals have been identified through genome-wide analyses in rice (Khan *et al*., 2019), *Medicago sativa* L. (alfalfa) (Wang *et al*., 2023), and in Tartary buckwheat (Ye *et al*., 2022). Finally, increasing genetic evidence links HIPP proteins to heavy metal homeostasis. In rice, loss of *HIPP42* gene function impairs tolerance to numerous heavy metal ions, such as Cd, Mn, Zn and Cu (Khan et al., 2019), while HIPP9 binds Cd and Cu, and its knock-out or overexpression significantly alters their accumulation (Xiong *et al*., 2023). Additionally, rice HIPP9 is proposed to facilitate Cu uptake in roots and contribute to Cd retention in nodes (Xiong et al., 2023).

Notably, HIPP proteins have also been shown to regulate phytohormone signaling. Clade I HIPPs have been identified as the regulators of cytokinin responses in Arabidopsis (Guo et al., 2021). They function as the components of the ER-associated protein degradation (ERAD) pathway, controlling the ERAD-dependent degradation of cytokinin-degrading CKX (cytokinin oxidase/dehydrogenase) proteins, and thereby cytokinin-dependent developmental processes (Guo et al., 2021). HIPPs are also involved in salicylic acid (SA) signaling. For example, Arabidopsis *HIPP3* is highly upregulated in response to *Pseudomonas syringae pv. tomato* infection, and its overexpression alters the expression of many genes associated with SA signaling (Zschiesche et al., 2015). Similarly, overexpression of *HIPP1-V* from *Haynalidia villosa* leads to increased transcript levels of SA-related genes, suggesting that HIPP1-V may be a positive regulator of SA signaling (Wang et al., 2023).

Despite recent advances in the characterization of *HIPP* genes, the biological functions of most members within the nearly 50-gene family in Arabidopsis remain largely unknown. In this study, we demonstrate that clade III HIPP proteins in Arabidopsis play a critical role in regulating plant growth and development. We show that these PD-localized HIPP proteins are essential for normal development and function across multiple stages of the plant life cycle. Our findings provide an important contribution towards understanding the function of the evolutionarily young HIPP protein family, expanding their functional scope beyond previously established roles in abiotic and biotic stress responses.

## RESULTS

### Clade III *HIPP* genes control vegetative shoot development

To investigate the physiological role of clade III HIPP proteins, we isolated single and higher-order loss-of-function mutants (Fig. S1) and analyzed their development. Two independent mutant alleles were isolated for each gene to ensure a robust assessment of gene function. *hipp* mutants exhibited altered rosette leaf development, with reduced rosette diameters in most mutants, particularly *hipp32* and *hipp32/33*, while *hipp34* and *hipp33/34* showed no significant changes (Fig. S2A-B). The leaf blade area was also significantly smaller, with the strongest, approx. 50%, reduction observed in *hipp32/33* and *hipp32/33/34* plants (Fig. 1A-B). Given that *hipp34*, *hipp32/34* and *hipp33/34* mutants showed no significant difference in leaf size (Fig. 1A-B), suggests that *HIPP32* and *HIPP33* are primary contributors to this trait. Morphologically, *hipp* mutants developed narrower, epinastic leaves with serrated margins (Fig. 1A, Fig. S2A). These phenotypic changes were more prominent in higher-order *hipp* mutants, suggesting functional redundancy among *HIPP* family members in regulating leaf morphology. Interestingly, *hipp* mutants also displayed a slight decrease in the number of formed rosette leaves (Fig. S2C). Collectively, these data suggest that clade III HIPP proteins are integral to the regulation of rosette and leaf development in Arabidopsis.

**Figure 1.**
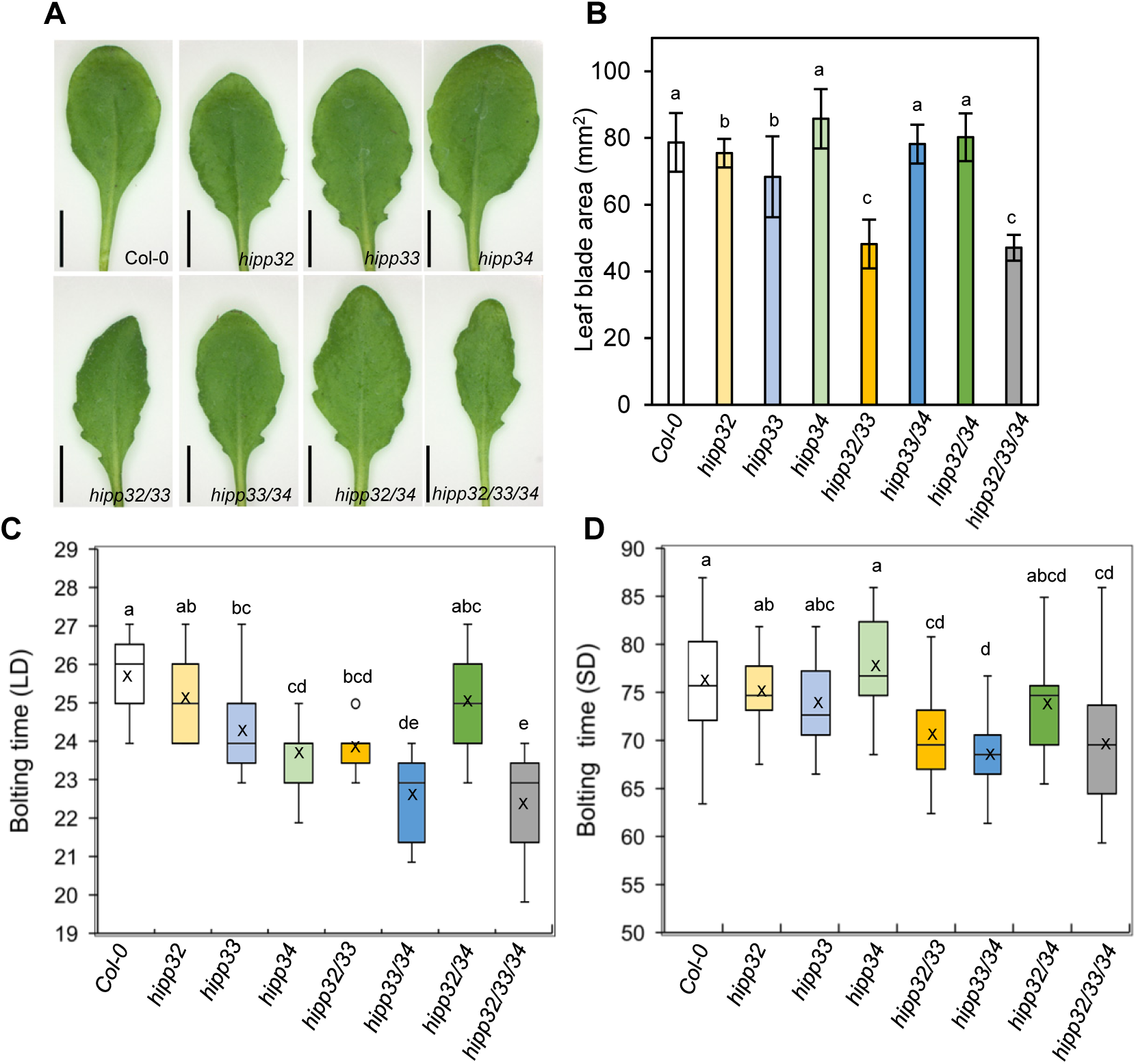
*hipp* mutants show abnormal rosette leaf development and early flowering. (A) Phenotypes of 7th rosette leave from wild-type (Col-0) and indicated *hipp* plants. Scale bars = 1 cm (B) Surface area of the 7th leaf at the bolting stage. Data are means ± SD (*n* ≥ 10). Flowering times of plants grown under (C) long-day (LD) and (D) short-day (SD) conditions. Data are means ± SD (*n* = 15). Different letters indicate individual groups for multiple comparisons with significant differences (one-way ANOVA, Wilcoxon, *p* < 0.05).

The primary inflorescence was significantly taller in *hipp33/34* and *hipp32/33/34* mutants as compared to the WT (Fig. S3A-B), suggesting that *HIPP33* and *HIPP34* act redundantly to negatively regulate shoot height. Most *hipp* mutants developed thinner stems, with the most pronounced reduction in *hipp32/33* and *hipp32/33/34* plants (Fig. S3C-D). Furthermore, clade III *HIPP* genes redundantly control shoot branching, as indicated by the increased axillary rosette branches in the *hipp33/34* and *hipp32/33/34* plants (Fig. S3E-F). Overall, these findings suggest that clade III HIPP proteins have largely overlapping functions in regulating multiple traits of shoot development in Arabidopsis.

### The roles of HIPP proteins in reproductive development

The *hipp* mutants exhibited an early flowering phenotype under standard long-day conditions (Fig. 1C). Specifically, *hipp33* and *hipp34* single mutants flowered up to 2 days earlier than the WT, and this phenotypic effect was significantly stronger in the *hipp33/34* and *hipp32/33/34* mutants (Fig. 1C), indicating that *HIPP33* and *HIPP34* function redundantly in repressing flowering time. The genetic analysis further suggests that HIPP32 has little to no influence in this process. Interestingly, the early flowering phenotype of *hipp* plants was less pronounced under short-day conditions (Fig. 1D), indicating that *HIPP* genes control flowering initiation in a light-dependent manner.

In addition, *hipp32/33*, *hipp33/34*, and *hipp32/33/34* mutants developed larger inflorescences compared to the WT, and increased silique formation rates were detected in these mutants (Fig. S4A-B), suggesting that the shoot meristem activity was enhanced. Total silique counts were also significantly higher in *hipp33/34* and *hipp32/33/34* mutants, yielding 30% and 40% more siliques, respectively, than WT (Fig. S4C). Moreover, these higher-order mutants remained reproductively active for an extended period (Fig. S4D), suggesting that the increased silique numbers were attributed to the enhanced meristematic activity and prolonged reproductive growth.

Higher-order *hipp* mutants also developed smaller flowers with partially altered morphology (Fig. 2A, Fig. S4E-F). *hipp32/33/34* often exhibited developmental defects such as fused organs and reduced stamen number (Fig. S4E). The reduction in flower size in *hipp* mutants was reflected by a corresponding decrease in silique size, which was particularly strongly affected in *hipp33/34* and *hipp32/33/34* (Fig. 2B). Seed production was significantly impaired, with *hipp33/34* siliques filled to 60% and *hipp32/33/34* rarely containing any seeds (Fig. 2C). The apparent ovule abortion (Fig. 2C) might partially result from reduced self-pollination due to reduced elongation of anther filaments (Fig. 2A), as well as reduced amounts of pollen in the mutants. To verify whether low fertility was due to female or male sterility, a series of controlled pollination tests were performed. Cross-pollination of *hipp33/34* and *hipp32/33/34* with WT pollen did not rescue the seed development and siliques elongation, whereas WT siliques developed normally when pollinated with *hipp33/34* or *hipp32/33/34* pollen (Fig. 2D), suggesting impaired female fertility in the mutants. These results underscore the critical role of *HIPPs* in female gametophyte development.

**Figure 2.**
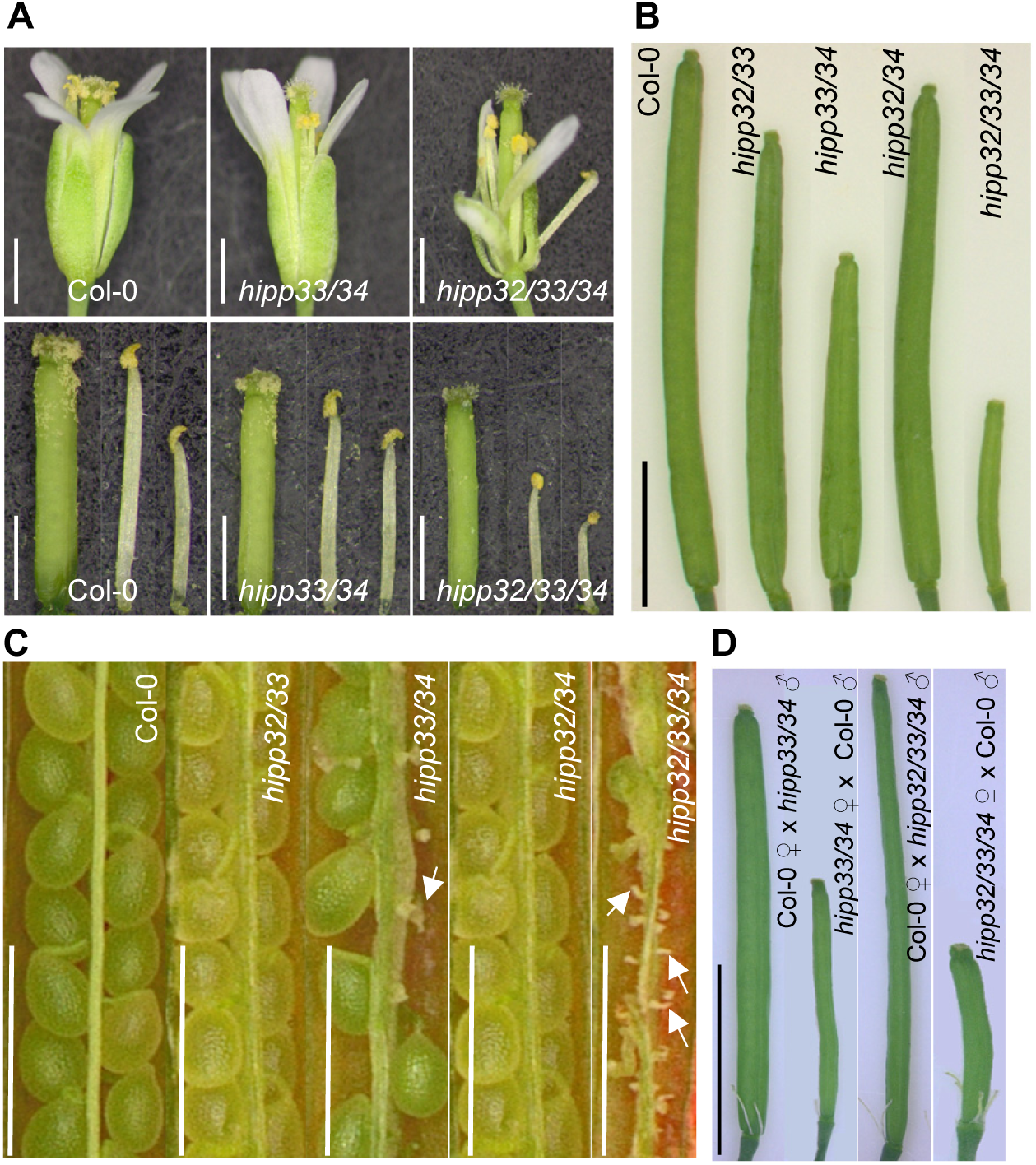
HIPP proteins control flower development and fertility in Arabidopsis. (A) Photos of flowers, stamens, and pistils from wild-type (Col-0) and the *hipp* plants at the pollination stage. Scale bars = 1 mm. (B) Fully developed siliques from the 5-week-old plants. Scale bar = 5 mm. (C) Seed filling in wild-type and the *hipp* plants. White arrows indicate aborted embryo sacs. Scale bar = 1 mm. (D) Siliques derived from crosses between wild-type and *hipp* plants. Scale bar = 5 mm.

### HIPPs are essential for shoot and root meristem development during embryogenesis

The low germination rate of *hipp32/33/34* seeds (20-50%, data not shown) prompted an analysis of potential defects during embryogenesis. Mature embryos of the *hipp32/33/34* mutant showed a range of developmental abnormalities, such as smaller and partially fused cotyledons (Fig. 3A-F), with some embryos developing three (Fig. 3C) or only one (Fig. 3F) cotyledon. Moreover, some *hipp32/33/34* embryos may undergo delayed or incomplete development, as their mature morphology resembled that of late torpedo-stage embryos (Fig. 3D). Compared to the WT embryos, the *hipp32/33/34* embryonic shoot apical meristem (SAM) was markedly enlarged (Fig. 3G-H), and, in some cases, the SAM layers and boundaries were less defined (Fig. 3I). Similarly, root apical meristem (RAM) establishment during embryogenesis was severely disrupted in *hipp32/33/34* (Fig. 3J-L). *hipp32/33/34* embryos showed regularly an abnormal cell patterning within the RAM with disorganized columella cells, and often indistinguishable quiescent center (QC) cells (Fig. 3J-L). Furthermore, unlike the wild type, *hipp32/33/34* embryos exhibited an accumulation of starch granules in the embryonic columella cells, indicating a premature differentiation in the RAM (Fig. 3J-L). Notably, an ectopic deposition of starch granules was also detected in other cells types within the embryonic root of *hipp32/33/34* mutants, suggesting that *HIPP* genes play a role in suppressing this process (Fig. 3L). Together, these findings highlight a critical function of clade III HIPPs in regulating embryonic patterning and development.

**Figure 3.**
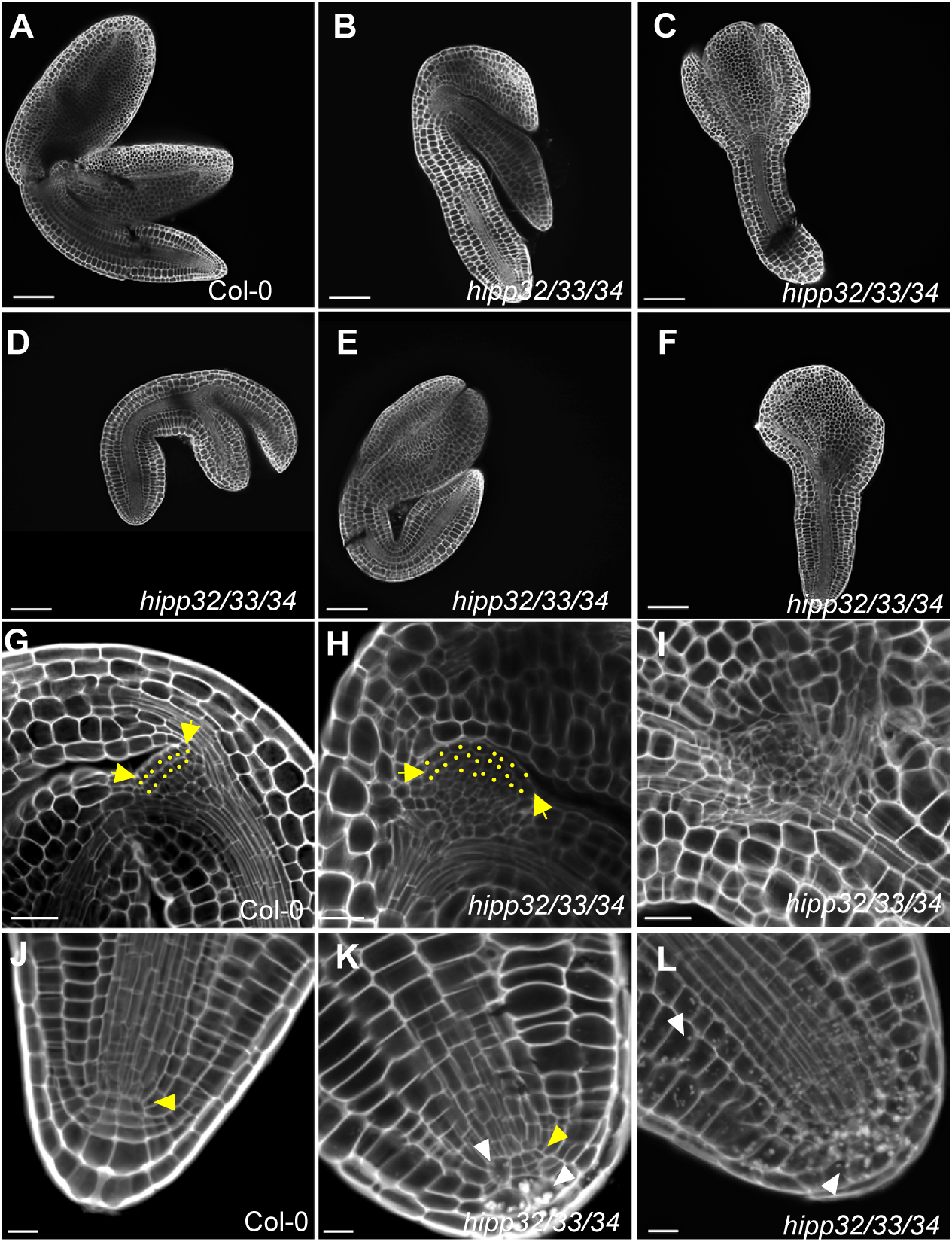
HIPP proteins regulate embryo development in Arabidopsis. (A-F) Mature embryos from wild-type (Col-0) and the *hipp32/33/34* plants visualized by pseudo-Schiff propidium iodide staining. Scale bars = 100 µm. (G-I) Images of shoot apical meristems (SAMs) from the wild-type (Col-0) and *hipp32/33/34* plants at the fully developed embryo stage. Arrows indicate the boundary between cotyledons and SAM. Yellow dots indicate individual cells in SAM. Scale bars = 25 µm. (J-L) Photos of root apical meristems (RAMs) from the wild-type (Col-0) and *hipp32/33/34* plants at the fully developed embryo stage. Yellow arrows indicate the quiescent center (QC) cells. White arrows indicate starch granules. Scale bars = 10 µm.

### HIPP proteins regulate root architecture and root apical meristem patterning

Root phenotypic analysis revealed that the primary root lengths of *hipp33* and *hipp34* were comparable to that of WT, whereas *hipp32* showed ∼30% reduction in primary root length (Fig. 4A). Interestingly, *hipp32/33* produced even shorter roots, while the loss of *HIPP34* abolished the short root phenotype in *hipp32* (Fig. 4A). These results suggest that *HIPP32* functions as a positive regulator of primary root growth and indicate a synergistic interaction with *HIPP33*, alongside an antagonistic relationship with *HIPP34* in regulating primary root growth. The lateral root (LR) number in *hipp32* was comparable to that of WT, while *hipp33* and *hipp34* mutants produced ∼30% more LRs (Fig. 4B). Higher LR number was also observed in *hipp32/34* and *hipp33/34* double mutants, but not in *hipp32/33*, indicating a complex genetic interaction among *HIPP* genes in regulating LR development. Lateral root density was elevated in all *hipps* (Fig. 4C), however, in the case of *hipp32* and *hipp32/33*, this increase was primarily attributed to a decrease in primary root lengths. The *hipp32/33/34* mutants exhibited severe root growth defects, with ∼20% germinated seedlings showing early primary root growth arrest. Remaining seedlings developed short primary roots with reduced gravitropism (Fig. 4D). In addition, *hipp32/33/34* roots had strongly disorganized RAM structure characterized by the absence of clearly defined QC cells, irregular cell division patterns, and frequent periclinal divisions in cortex and epidermal cells (Fig. 4E). Importantly, the effects of individual *hipp* mutations on root development were fully recapitulated using independent loss-of-function alleles (Fig. S5A-C), confirming the causal relationship and supporting the conclusion about the essential role of HIPPs in regulating root system architecture and meristem patterning.

**Figure 4.**
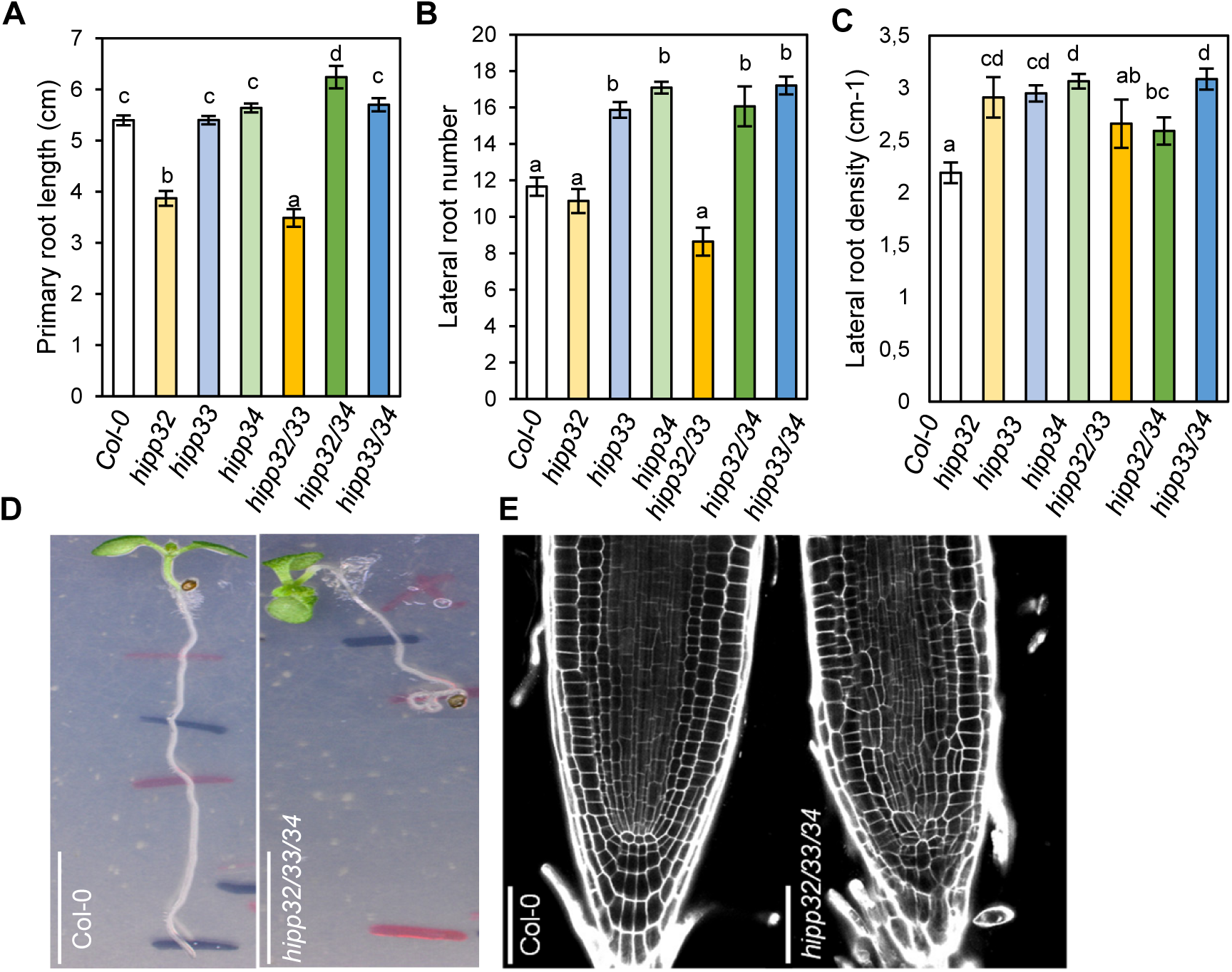
HIPP proteins control root system architecture in Arabidopsis. Quantification of the (A) primary root length, (B) lateral root number, and (C) later root density in the 11-day-old wild-type (Col-0) and *hipp* plants. Data are means ± SE (*n* ≥ 10). Different letters indicate individual groups for multiple comparisons with significant differences (one-way ANOVA, Wilcoxon, *p* < 0.05). (D) Photos of primary roots from the wild-type and *hipp32/33/34* plants. Scale bars = 50 mm. (E) Confocal images of root apical meristems from 6-day-old plants. Scale bars = 10 µm.

### HIPPs influence cytokinin responses without major changes in cytokinin hormone levels

Given the detected root phenotypic alterations and previously reported role of clade I HIPPs in regulating cytokinin responses (Guo et al., 2021), we explored the possible link between the analyzed *HIPP* genes and cytokinin signaling by analyzing the cytokinin reporter *TCSn:GFP* (Zürcher *et al*., 2013) introgressed into *hipp* mutants. Visualization of root meristems revealed an overall reduction of *TCSn:GFP* signal in *hipp* plants, with the strongest reduction detected in *hipp33* and *hipp33/34* (Fig. 5A-B). *HIPP32* had only weak effect on *TCSn:GFP* activity, apparently acting antagonistically to *HIPP33* and *HIPP34* in modulating cytokinin output (Fig. 5A-B). Given that cytokinin signaling inhibits root growth (Werner *et al*., 2003), it is noteworthy that the short-root phenotype of *hipp32* and *hipp32/33* was not correlated with increased *TCSn:GFP* activity.

**Figure 5.**
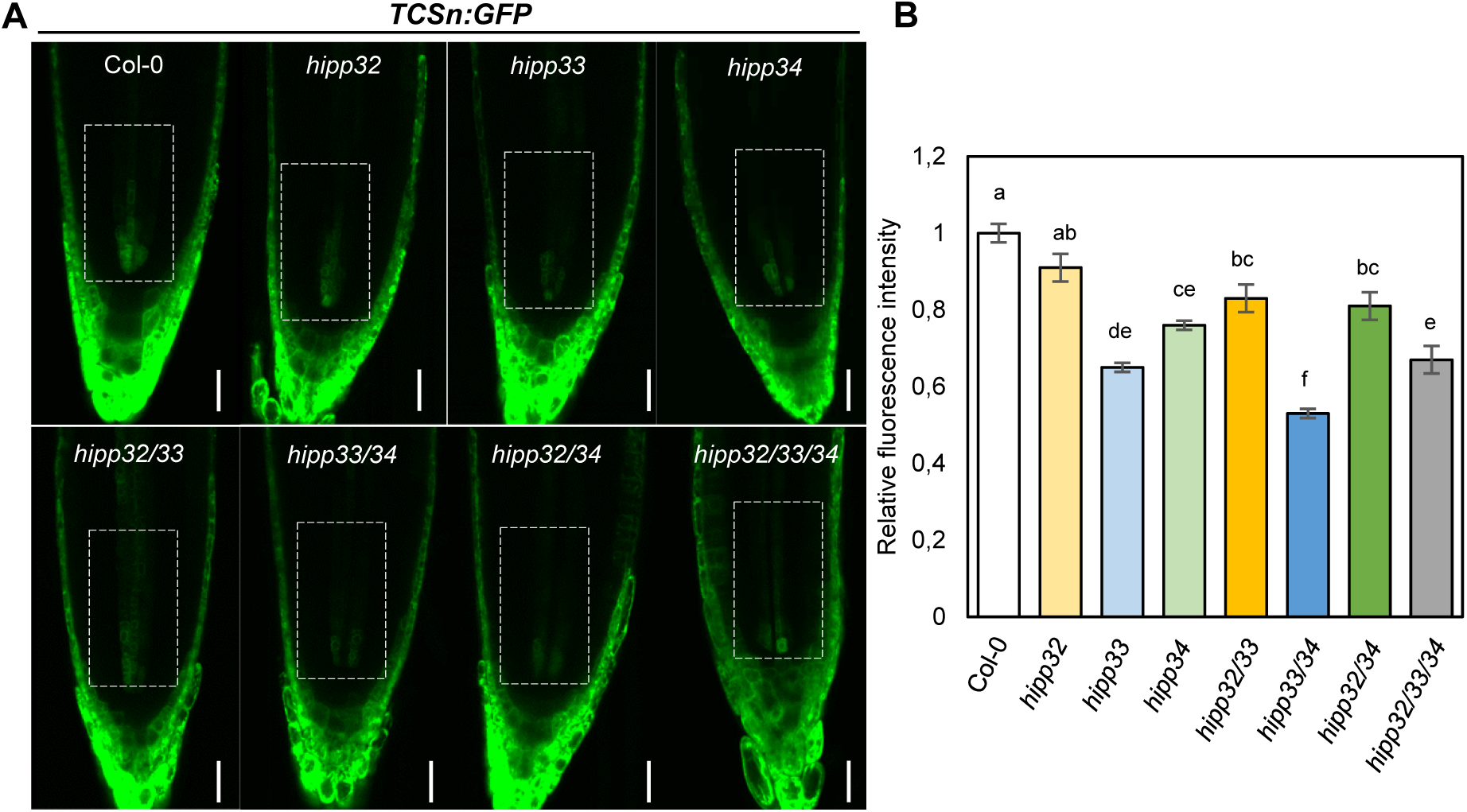
*hipp*s show attenuated cytokinin signaling. (A) Confocal images of the *TCSn:GFP* reporter gene in roots of the wild-type (Col-0) and *hipp* plants. Scale bars = 25 μm (B) Quantification of the *TCSn:GFP* fluorescence in white boxes in A. Data are means ± SE (*n* ≥ 10). Different letters indicate individual groups for multiple comparisons with significant differences (one-way ANOVA, Wilcoxon, *p* < 0.05).

Interestingly, the *hipp32* and *hipp33* plants were more sensitive to exogenously applied cytokinin, whereas *hipp34* showed a response comparable to WT (Fig. S6A-B). Enhanced CK sensitivity was also observed in *hipp32/33* and *hipp33/34* (Fig. S6A-B). Further increase in cytokinin sensitivity in *hipp32/33/34* may suggest that clade III HIPP proteins function cooperatively to modulate plant sensitivity to CK.

Cytokinin profiling in *hipp* mutants revealed that, despite reduced *TCSn*:*GFP* activity, the overall cytokinin content remained largely unchanged (Fig. S7). The most significant changes were the reduced levels of cytokinin free bases and ribosides in *hipp32/33* mutants compared to WT (Fig. S7, Table S1). Taken together, the results indicate that *HIPP* genes can modulate cytokinin responses, however the changes in apparent cytokinin signaling do not clearly explained the phenotypic alterations observed in the mutants, suggesting that the cytokinin-related effects might be secondary, at least with regard to root meristem activity.

### Transcriptomic profiling reveals a role for clade III *HIPP*s in regulating auxin responses

To further explore molecular pathways through which clade III HIPPs regulate plant development, we conducted transcriptomic profiling of 5-day-old WT and *hipp* single mutant seedlings. Among the genotypes, *hipp32* showed the largest number of differentially expressed genes (DEGs) (843), followed by *hipp33* (280) and *hipp34* (183) compared to WT (Fig. 6A). A significant overlap of DEGs was observed among individual *hipp* mutants, supporting the notion of high functional redundancy within the clade III HIPP family. This overlap was particularly pronounced in *hipp33* and *hipp34*, with only 12% and 20% of DEGs unique to each genotype, respectively (Fig. 6A). In contrast, 68% of the DEGs identified in *hipp32* were unique, suggesting that while *HIPP32* shares functional redundancy with *HIPP33* and *HIPP34*, it also regulates a distinct set of genes (Fig. 6A). The Gene Ontology (GO) analysis of the 99 DEGs deregulated across all *hipp* mutants revealed a significant enrichment for GO terms related to the plant hormone auxin (“response to auxin” and “auxin-activated signaling”) (Fig. 6B). Similarly, the Kyoto Encyclopedia of Genes and Genomes (KEGG) pathway enrichment analysis indicated that these 99 DEGs were primarily associated with the “plant hormone signal transduction” pathway (Fig. 6C). Among the 10 genes associated with this KEGG pathway, 9 were related to auxin, while only one gene (*Arabidopsis response regulator* 6; *ARR6*) was linked to cytokinin (Fig. 6D). Interestingly, all auxin-associated genes were down-regulated in *hipp* mutants, whereas *ARR6* was upregulated relative to WT (Fig. 6D). The GO and KEGG enrichment analyses of DEGs from individual *hipp* mutants further supported the potential role of HIPPs in modulating auxin signaling, and identified additional pathways regulated by these proteins (Fig. S8-9). Together, these findings suggest that clade III HIPPs primarily modulate auxin-related transcriptional programs, with a relatively minor role in regulating cytokinin responses.

**Figure 6.**
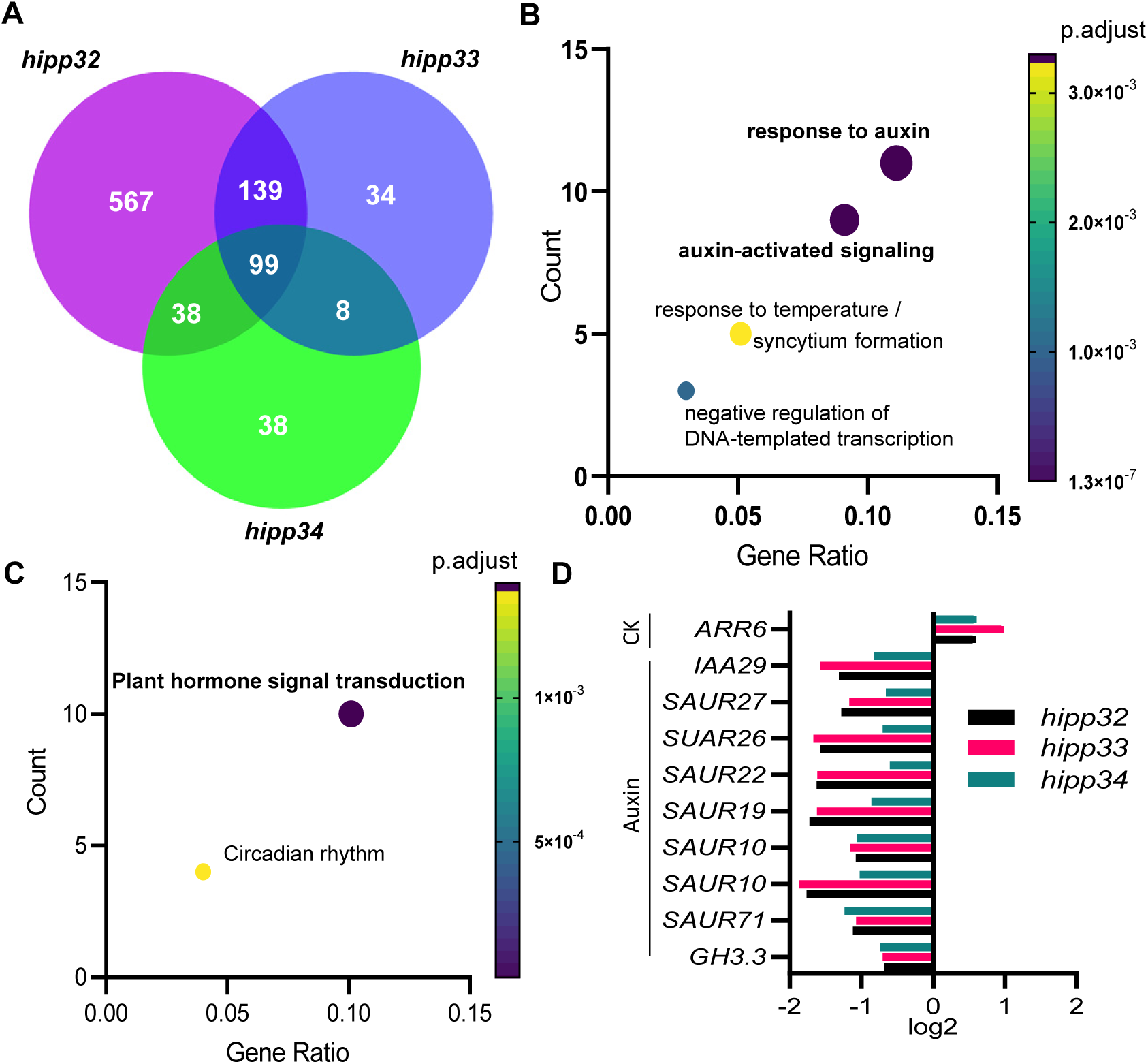
HIPP proteins regulate auxin responsiveness and signaling. (A) Venn diagram showing differentially regulated genes in *hipp32*, *hipp33*, and *hipp34*. (B) Top 5 enriched GO terms of the 99 deregulated genes detected in all *hipp* mutants. (C) Two enriched KEGG pathways of the 99 deregulated genes in the *hipp* plants. (D) Diagram showing expression (log2 fold change) of all 10 genes belonging to the ‘’Plant hormone signal transduction’’ KEGG category in (C). CK, cytokinin.

### Clade III HIPP proteins are specifically targeted to plasmodesmata

To better understand the molecular mode of action of the analyzed HIPPs, we examined their subcellular localization. The proteins were fused to GFP at their N termini to prevent interference with the potential C-terminal prenylation, and the fusion proteins were expressed in *N. benthamiana* leaf epidermis, using the constitutive CaMV 35S promoter. Confocal microscopy revealed that GFP-HIPP33 and GFP-HIPP34 exhibited highly specific localization to discrete punctate structures at the cell periphery (Fig. 7B-C), consistent with a plasmodesmal (PD) distribution pattern (Oparka *et al*., 1997). GFP-HIPP32 displayed a comparable punctate peripheral localization, although the signal was less distinctly defined (Fig. 7A).

**Figure 7.**
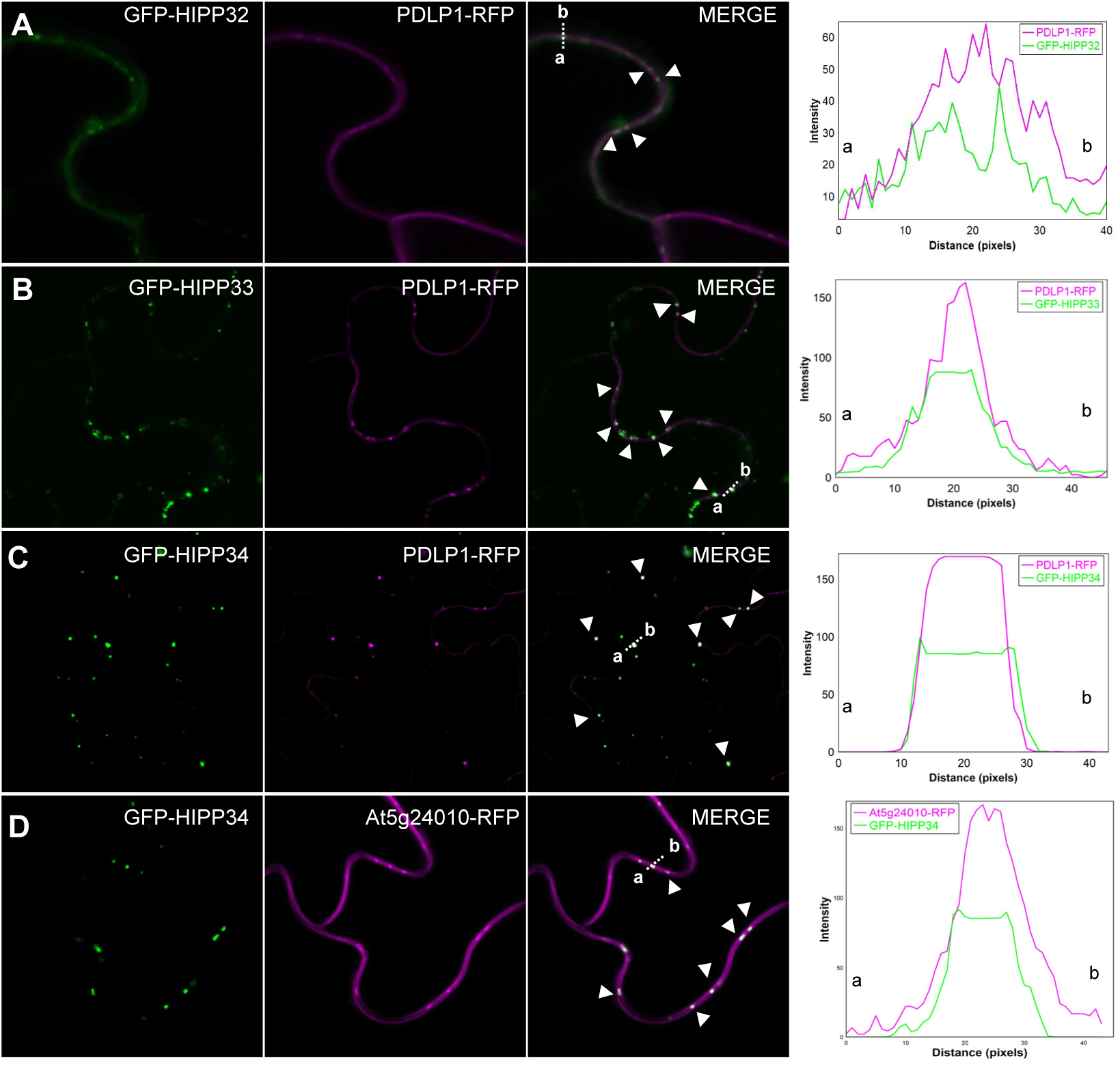
Clade III HIPP proteins localize to plasmodesmata. Subcellular localization of (A) GFP-HIPP32, (B) GFP-HIPP33, and (C) GFP-HIPP34 co-expressed transiently with the plasmodesmata marker PDLP1-RFP in *N. benthamiana* leaves. (D) Subcellular localization of GFP-HIPP34 in leaf epidermis cells of stably transformed *35S:GFP-HIPP34* Arabidopsis plants crossed with the plasmodesmata marker line At5g24010-RFP. White arrowheads indicate regions of HIPP localization at the PD. Dotted lines in the merge panels correspond to the distance and direction of the line intensity plot.

To assess the potential localization to PD, HIPP fusion proteins were co-expressed with the PD marker plasmodesmata-located protein 1 (PDLP1;Amari *et al*., 2010). All three GFP-HIPP proteins showed a colocalization with PDLP1-RFP, supporting their localization at PD (Fig. 7A-C). To validate the PD localization of HIPPs, Arabidopsis plants stably expressing the *35S:GFP-HIPP34* transgene were generated and crossed with the reporter line expressing an RFP-tagged CrRLK-like protein, which is enriched at PD and has been previously used as a PD marker (35S:At5g24010-RFP; Fernandez-Calvino *et al*., 2011, Oulehlová et al., 2019). Confocal microscopy of homozygous progenies confirmed the co-localization of the two proteins (Fig. 7D) and confirmed the GFP-HIPP34 residency at PD.

To understand the mechanism of HIPP targeting to PD, we analyzed potential protein targeting domains. To test whether prenylation of clade III HIPPs is relevant for PD localization complex formation, we mutated the prenyl-accepting Cys residue within the isoprenylation motif of HIPP34 (HIPP34^C470G^) and fused it to GFP. We also generated a HIPP34 mutant version lacking the functional HMA domain by mutating the metal-binding Cys residues (HIPP34^hma^). Confocal microscopy showed that the frequency of PD localization was lower and the fluorescence signal associated with PD was significantly weaker in *N. benthamiana* plants expressing GFP-HIPP34^C470G^ (Fig. S10), suggesting that prenylation is largely required for the localization of HIPP34 to PD. Similar reduction in PD-specific localization was observed also for the GFP-HIPP34^hma^ fusion protein (Fig. S10), indicating that HMA domain is relevant for the targeting to PD as well.

### Symplasmic transport is altered in *hipp* mutants

PD are intercellular channels across the plant cell walls that mediate and facilitate the symplasmic molecular transport between neighboring cells, which is required for plant growth and development (Benitez-Alfonso, 2014). The symplasmic transport relies on the permeability of PD and several PD-localized proteins have been shown to regulate the process (Sager & Lee, 2018). Given the observed localization of HIPP proteins to PD, we investigate their possible function at PD by analyzing the symplasmic permeability in *hipp* mutants using the Arabidopsis plants expressing GFP from the phloem companion cell-specific *Sucrose-H^+^ Symporter 2* gene promoter (*pSUC2:GFP*) (Imlau *et al*., 1999). The expressed GFP moves from the companion cells into sieve elements and is symplastically unloaded via PD from phloem cells into sink tissues. As previously shown by others, the GFP fluorescence in the sink tissues, such as the root meristem, can serve as an indirect measure for cellular connectivity (Vatén *et al*., 2011, Benitez-Alfonso *et al*., 2009).

Confocal microscopy revealed increased GFP intensity in the root meristems of *hipp32*, *hipp34*, *hipp33/34*, and *hipp32/34* mutants compared to the wild type (Fig. 8A). To assess the GFP movement from the phloem into the meristematic region, the ratio of GFP fluorescence in the proximal meristem to a defined region in the vascular cylinder above transition zone was determined. The ratios were significantly elevated in *hipp33* and *hipp34* (Fig. 8A-B), indicating enhanced GFP movement in the mutant roots. An additive effect was observed in the double mutant *hipp33/34*, where GFP movement was increased by 25% relative to the wild type (Fig. 8B). While *hipp32/34* showed a trend toward higher values than *hipp34*, this difference was not statistically significant (Fig. 8B). No significant changes were detected in *hipp32* and *hipp32/*33 (Fig. 8B). These results collectively suggest that *HIPP33* and *HIPP34* function redundantly to restrict PD permeability and cellular connectivity in root tissues.

**Figure 8.**
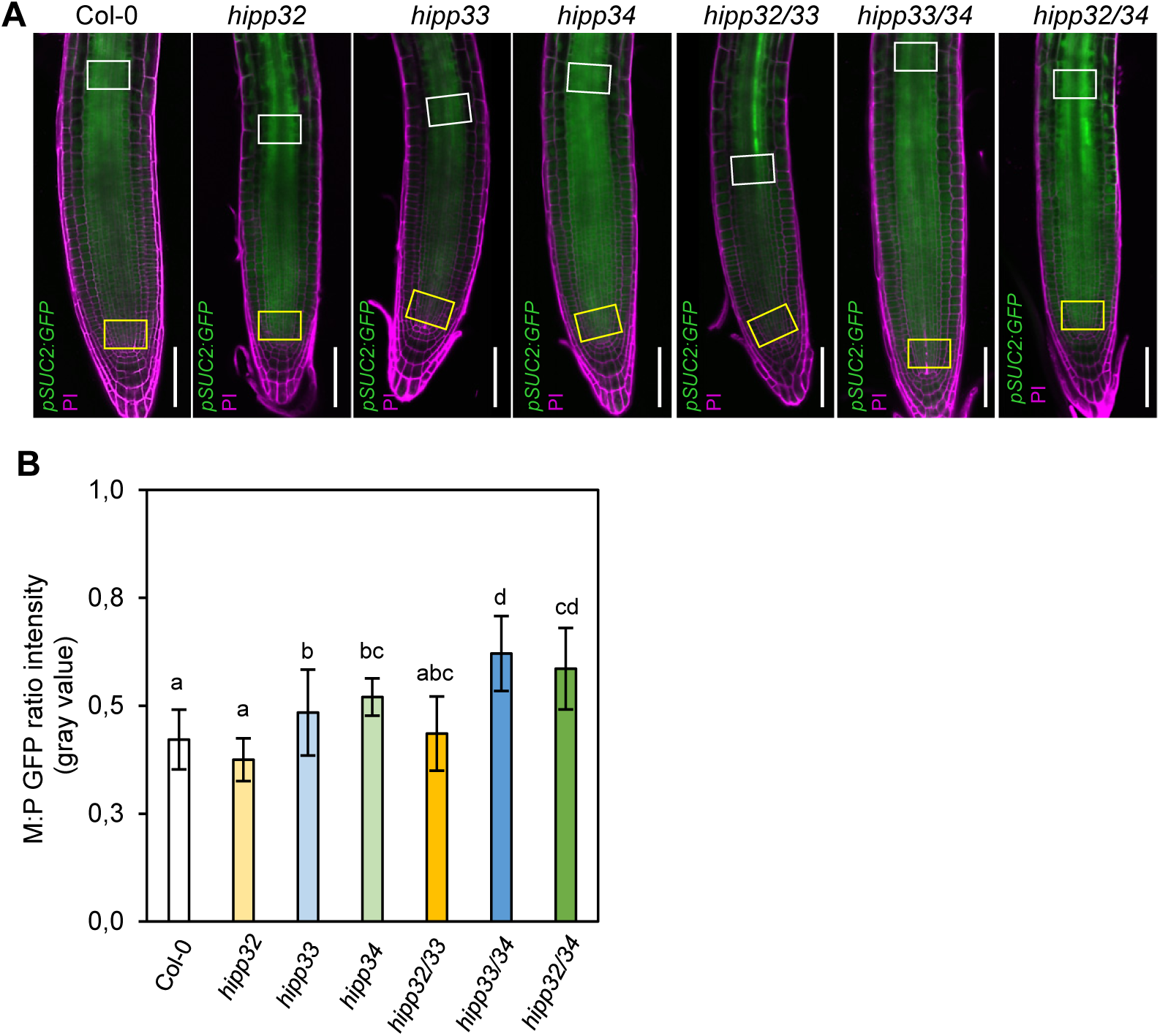
*HIPP33* and *HIPP34* regulate phloem unloading into the root meristem. (A) Confocal images of the 5-day-old primary root of wild-type (Col-0), *hipp* mutants expressing the mobile phloem marker *pSUC2:GFP (*green). Propidium iodide (PI, magenta) was used for cell visualization. Scale bars = 50 μm (B) GFP movement was determined as the ratio of GFP mean fluorescence intensity in the root meristem above the QC (yellow box) relative to the phloem/vascular cylinder area above the transition zone (white box). Data are means ± SD (n = 6-24). Different letters indicate individual groups for multiple comparisons with significant differences (one-way ANOVA, Wilcoxon, p < 0.05).

## DISCUSSION

HIPP proteins, defined by the presence of the HMA domain and posttranslational prenylation, have uniquely evolved in vascular plants. While their functional roles remain largely unexplored, emerging evidence suggests their involvement in heavy metal homeostasis, abiotic stress regulation, and plant immunity (reviewed by Barr et al., 2023, Turley & Faulkner, 2025). In comparison, their role in shaping plant development has been studied to a much lesser extent. This study reveals that the distinct phylogenetic subgroup III of HIPP genes plays a regulatory role in various aspects of plant growth, as reflected by the pleiotropic phenotypic defects observed in the corresponding mutant lines.

The wide-ranging roles of clade III HIPP genes in plant development are consistent with their diverse expression patterns observed throughout plant development (Fig. S11-13). The analysis of transgenic Arabidopsis plants expressing individual *pHIPP:GUS* reporter constructs demonstrated that the corresponding genes are expressed in several different, largely spatially restricted domains (Fig. S11-13). Interestingly, all three genes show overlap in some tissues and organs, such as roots and young leaves, which correlates with the complex genetic interactions between the genes revealed by the mutant phenotypic analyses. These overlapping expression patterns likely contribute to the functional redundancy observed among clade III HIPP genes, while their distinct expression domains may underlie their unique roles in specific developmental processes.

The RNA-seq data support the notion that clade III HIPP proteins exhibit a high degree of functional redundancy in controlling developmental processes, as evidenced by the significant overlap in differentially expressed genes (DEGs) among the corresponding single mutants. This redundancy is consistent with phenotypic analyses, where higher-order mutants often display more pronounced developmental phenotypes. For example, the triple mutant *hipp32/33/34* exhibited severe defects in embryogenesis, shoot and root meristem development, and reproductive growth. Functional redundancy is particularly evident for HIPP33 and HIPP34, with only 12% and 20% of DEGs being unique to *hipp33* and *hipp34*, respectively. In contrast, while HIPP32 also shares targets with HIPP33 and HIPP34, it regulates a substantially larger proportion of unique genes – 68% of its DEGs are not shared – suggesting additional functions beyond those shared with HIPP33 and HIPP34. These findings align with the calculated protein homology within clade III members, where HIPP33 and HIPP34 share the highest sequence identity (67%). Additionally, the eventual unique regulatory role of HIPP32 is further supported by its partially distinct subcellular localization, compared to HIPP33 and HIPP34.

KEGG pathway analysis of DEGs shared among the *hipp32*, *hipp33*, and *hipp34* mutants, as well as DEGs from single *hipp* mutants, revealed the role of HIPP proteins in regulating plant hormone signaling pathways. This regulatory function likely contributes to the diverse developmental defects observed in *hipp* mutants. Notably, GO enrichment analysis identified a significant role of HIPP proteins in auxin regulation, as demonstrated by the enrichment of terms such as “response to auxin” and “auxin-activated signaling pathway” among the top five enriched GO terms. These findings highlight the critical role of HIPPs in auxin-related transcriptional programs. Interestingly, while clade III HIPPs were previously detected among proteins interacting with cytokinin oxidase/dehydrogenase (CKX) enzymes in yeast-based experiments (Guo et al., 2021), the transcriptomic data suggest that their connection to cytokinin signaling might be limited. Among the DEGs associated with the KEGG plant hormone signal transduction pathway, only a single gene, *ARR6*, is linked to cytokinin, while the remaining genes are associated with auxin signaling. Moreover, no notable changes in cytokinin levels were detected in the *hipp* mutants, reinforcing the notion that HIPPs play only a minor role in cytokinin regulation. Alternatively, their functional connection to cytokinin activity may be restricted to particular developmental stages or processes that were not captured in this study, necessitating more experimental investigation in the future. In this line, it is worth noticing that the *TCSn:GFP* reporter activity was altered in *hipp* mutant roots.

An important finding of this study is the specific targeting of clade III proteins to PD. Interestingly, several previous reports have shown PD localization of other HIPP proteins outside of clade III, such as HIPP1 and HIPP7 in Arabidopsis (Guo et al., 2021), HIPP26 in *N. benthamiana* (Cowan et al., 2018), and HIPP43 in barley (Were et al., 2025). Interestingly, these previously reported HIPP proteins exhibited additional subcellular localizations besides PD, such as the endoplasmic reticulum (ER) (Guo et al., 2021) and the nucleus (Cowan et al., 2018). In contrast, GFP-HIPP33 and GFP-HIPP34 proteins displayed apparently specific localization at the PD. PD localization of HIPP proteins is further supported by proteomic studies, where several HIPP proteins, including clade III members, have been identified as components of the PD proteome (Brault *et al*., 2019, Kraner *et al*., 2017, Johnston et al., 2023). The mechanisms underlying protein targeting to PD remain largely unknown. To date, no universal sequence or mechanism has yet been found for targeting proteins to plasmodesmata. Our data indicate that the isoprenylation motif and the HMA domain contribute to PD targeting of clade III HIPP proteins. The involvement of isoprenylation in PD targeting has been previously described for HIPP7 (Guo et al., 2021), HIPP26 (Cowan et al., 2018) and HIPP43 (Were et al., 2025). Interestingly, another posttranslational modification, S-acylation, has also been shown to play a role in PD targeting (Cowan et al., 2018). It would be interesting to analyze whether clade III HIPPs also undergo S-acylation, and whether this modification contributes to their localization to PD.

Collectively, an increasing line of evidence suggests that PD localization may be a conserved and likely functionally relevant characteristic of many HIPP proteins. However, the functional relevance of this PD localization is currently unclear. PD-localized HIPP26 (Cowan et al., 2018) and HIPP43 (Were et al., 2025) are targeted by pathogens, and given that plasmodesmata facilitate intercellular pathogen and effector spread, PD-associated HIPPs may act as endogenous regulators of PD aperture and thus represent targets for pathogens to maintain cell-to-cell connectivity during infection (reviewed by Turley & Faulkner, 2025). Furthermore, as this work clearly establishes a role for clade III HIPPs in regulating diverse aspects of plant development – including embryogenesis, maintenance of apical meristems, root architecture, shoot branching, leaf morphogenesis, and floral organ formation it is tempting to speculate that HIPP proteins display their activity by regulating PD function. Analysis of symplastic transport in *hipp* mutants using the *pSUC2:GFP* reporter supports this hypothesis, demonstrating that HIPP33 and HIPP34 act redundantly to restrict PD permeability and limit cellular connectivity in root tissues. It would be interesting to address the question of whether the altered hormonal responses observed in *hipp* mutants might also be linked to altered PD activity. In the context of the described changes in hormonal activity in *hipp* mutants, it is interesting that PD-mediated transport has recently been discussed as an important mechanism of hormone distribution between cells (reviewed by Zanini & Burch-Smith, 2024). For example, besides the specific protein-based transport mechanism on the membrane, cell-to-cell trafficking of auxin via PD has been recognized as important for establishing auxin gradients (Gao *et al*., 2020, Mellor *et al*., 2020, Band, 2021). In particular, PD-dependent auxin transport has been implicated in lateral root development (Sager *et al*., 2020), regulation of cell division in the root apical meristem (Olatunji *et al*., 2023), and suppressed lateral root formation in response to transient water stress (Mehra *et al*., 2022).

In conclusion, clade III HIPP proteins represent a unique group of PD-targeted regulators with pleiotropic roles in plant development. Our genetic and molecular analyses demonstrate that clade III *HIPP* genes exhibit both overlapping and distinct roles across multiple developmental processes. *HIPP33* and *HIPP34* function redundantly in regulating shoot architecture, reproductive traits, and plasmodesmata permeability, whereas *HIPP32* uniquely contributes to primary root growth and modulates a distinct set of genes. This combinatorial activity underlies the pleiotropic phenotypes observed in *hipp* mutants and highlights the complexity of HIPP-mediated developmental control. The ability of HIPPs to modulate auxin signaling and PD activity underscores their importance in shaping plant growth and architecture. Therefore, elucidating the functional role of HIPP proteins at PD, particularly how the PD localization of clade III HIPPs contributes to auxin signaling regulation, remains an important area for future research.

## MATERIALS AND METHODS

### Generation of gene constructs for expression in plants

Genomic sequences of *HIPP32* and *HIPP33* were amplified from Arabidopsis Col-0 genomic DNA. cDNA of HIPP34 was amplified from the SALK clone U09068 / U13015. Primers used are listed in Table S2. The fragments were cloned into pDONR221 (Invitrogen) and subsequently subcloned by Gateway LR recombination into pB7FWG2 or pK7WGF2, to generate N-terminal GFP fusion constructs under the control of the cauliflower mosaic virus (CaMV) 35S promoter. The HIPP34^C470G^ and HIPP34^hma^ mutations were introduced by the QuikChange site-directed mutagenesis kit (Stratagene) and subcloned from pDONR221 into pK7FWG2. For the CRISPR/Cas9-mediated targeted editing of *HIPP32* and *HIPP33*, the site-specific sgRNA oligonucleotides were cloned into pHEE401E and transformed into *hipp34-2* and Col-0 WT Arabidopsis, respectively, by *Agrobacterium tumefaciens* (GV3101:pMP90) mediated floral dip method. To construct β-glucuronidase (GUS) reporter fusions clones, a 2.1 kb upstream region of *HIPP32,* a 2.2 kb upstream region of *HIPP33* and 2 kb upstream region of *HIPP34* were amplified by PCR using primers listed in Table S2. The amplified promoters of *HIPP32* and *HIPP33* were cloned into pDONR222 (Invitrogen), and subsequently subcloned into pBGWFS7 vector using Gateway LR recombination (Invitrogen), while the *HIPP34* promoter was cloned upstream of the *GUS* gene into the pCB308 vector.

### Plant material and growth conditions

*Arabidopsis thaliana* ecotype Columbia-0 (Col-0) was used as the wild type. The following lines were described previously: *TCSn:GFP* (Zürcher & Müller, 2016), *pSUC2:GFP* (Imlau et al., 1999), and *35S:At5g24010-RFP* (Fernandez-Calvino et al., 2011). The T-DNA insertion mutants *hipp32-1* (SALK_017337), *hipp32-2* (SAIL_273_B10), *hipp33-1* (SAIL_1235_G02), *hipp33-2* (SAIL_899_D10C1), *hipp34-1* (SALK_079319C), *hipp34-2* (WiscDsLox248H10) were obtained from the Nottingham Arabidopsis Stock Centre. Double *hipp32,33* (*hipp32-2*, *hipp33-2*), *hipp32,34* (*hipp32-3*, *hipp34-2*), *hipp33,34* (*hipp33-2*, *hipp34-2*), and triple *hipp32,33,34* (*hipp32-3*, *hipp33-2*, *hipp34-2*) mutants were generated by genetic crosses and CRISPR/Cas9-mediated mutagenesis.

Arabidopsis plants were grown *in vitro* on half-strength Murashige and Skoog (MS) medium containing 1 g/L sucrose or in the green house under long-day conditions (16h light / 8h dark; 21 / 18°C). *N. benthamiana* plants were grown at 24°C under 14 h light/10 h dark conditions in the greenhouse.

### Transient expression in *N. benthamiana*

Gene constructs were transformed as described previously (Niemann *et al*., 2018, Sparkes *et al*., 2006) using *A. tumefaciens* strain GV3101:pMP90 and 6-week-old *N. benthamiana* plants.

For co-expression, the Agrobacterium cultures were mixed in infiltration medium to a final OD_600_ of 0.1 for each construct. *35S:p19 (*Voinnet *et al*., 2003) was included in all infiltrations at OD_600_ 0.1. 35S:PDLP1-RFP (Amari et al., 2010) was used as a PD marker. The fluorescence signal was analyzed 48-72 h after infiltration.

### Microscopy

Confocal laser scanning microscopy analysis was performed using Leica TCS SP5 and Stellaris 5 microscopes. For subcellular analysis, the GFP- and RFP-derived fluorescence was detected using excitation wavelengths of 488 nm and 561 nm, respectively, with corresponding emission wavelengths at 498-538 nm for GFP and 590-650 nm for RFP. To visualize cell organization in roots, roots were stained with 0.1 mg/ml propidium iodide (PI).

Arabidopsis embryos were analyzed by the modified pseudo-Schiff propidium iodide (mPS-PI) staining technique (Truernit *et al*., 2008). Briefly, embryos were fixed in 50% methanol and 10% acetic acid at 48°C for at least 12 h, then transferred to 80% ethanol and incubated at 80°C for 5 min. After an additional 1 h in fixative, roots were rinsed with water, incubated in 1% periodic acid at room temperature for 40 min and stained with Schiff reagent containing propidium iodide for 2 h. Samples were mounted in chloral hydrate solution at 4°C overnight before confocal microscopy analysis.

For the *HIPP* promoter activity studies, histochemical analysis of the GUS reporter enzyme was performed as described previously (Werner et al., 2003), and microscopic analysis was carried out using Axioskop2 plus microscope (Zeiss) and a light stereomicroscope (SZX12, Olympus).

### Hormone and chemical treatments

For cytokinin treatment, Arabidopsis seedlings were grown on half-strength MS medium for 5 days, and transferred to half-strength MS liquid medium supplemented with BA (1 μM) or mock (0.001% DMSO) for 16 h prior to sample collection.

### Cytokinin measurements

The cytokinin contents in shoots of soil-grown plants at different developmental stages were determined by ultra-performance liquid chromatography-electrospray-tandem mass spectrometry as described by (Svačinová *et al*., 2012), including modifications described by (Antoniadi *et al*., 2015).

### RNA-Seq

For RNA-seq analysis, seedlings were grown on half-strength MS media for 5 days. Three biological replicates per genotype were collected, each consisting of 10 seedlings. Total RNA was extracted using NucleoSpin RNA (Machery & Nagel) extraction kit according to the manufacturer’s instructions. BGI Genomics (Hong Kong, China) performed mRNA enrichment and library preparation. The libraries were sequenced using the DNBSEQ-G50 platform. The sequencing generated in total approx. 387 million reads. A total of 372,099,461 reads that passed the quality filter were mapped to the TAIR10/Araport11 *A.thaliana* Col-0 genome assembly using HISAT2. DEGs were selected based on the adjusted *P*-value (FDR < 0.05) and 1.5-fold change. The Gene Ontology (GO) enrichment and the Kyoto Encyclopedia of Genes and Genomes (KEGG) pathway analysis were conducted using DAVID 6.8 software (Huang *et al*., 2009, Sherman *et al*., 2022).

## ACCESSION NUMBERS

Sequence data from this article can be found in the GenBank/EMBL libraries under the following accession numbers: *HIPP32* (At3g05220), *HIPP33* (At5g19090), *HIPP34* (At3g06130).

## ACKNOWLEDGEMENTS

This work was supported by grants from the Austrian Science Fund (M 2815-B to A.M.G., and P 30945 to T.W.), and by a doctoral fellowship from the Austrian Academy of Sciences to C.B.A.

## AUTHOR CONTRIBUTIONS

G.L., A.M.G., and T.W. planned and designed the research. G.L., C.B.A., I.B., H.W., A.M.G, O.N., and T.W. performed the research and analyzed the data. A.M.G. and T.W. wrote the manuscript.

---

**Figure S1.** Molecular characterization of clade III *HIPP* gene loss-of-function alleles. (A) Gene structure and positions of the T-DNA insertions and CRISPR-Cas9-induced mutations in the *hipp* mutants. Triangles indicate the positions of the T-DNA insertions. PAM sequences adjacent to the sgRNA target sites are underlined. Base pairs marked in red indicate editing events in the CRISPR-Cas9 mutants. The CRISPR-Cas9 editing caused frameshift mutations, resulting in premature stop codons in *HIPP32* and *HIPP33* genes. (B) *HIPP32*, *HIPP33* and *HIPP34* transcript levels in wild-type (WT) and T-DNA insertion *hipp* mutants. Semi-quantitative RT-PCR analysis using primers flanking the T-DNA insertions (Table S2) did not detect any transcripts in the respective mutant lines, indicating that they are null alleles. *ACTIN7* was used as a control.

**Figure S2.** Loss of *HIPP* gene function leads to abnormal rosette leaf development. (A) Photos of rosettes from the 24-day-old wild-type (Col-0) and *hipp* plants. (B) Rosette diameters and (C) rosette leaf numbers of plants shown in (A). Data are means ± SD (*n* ≥ 10). Different letters indicate individual groups for multiple comparisons with significant differences (one-way ANOVA, Wilcoxon, *p* < 0.05).

**Figure S3**. *hipp* mutants show altered shoot growth and increased inflorescence meristem activity. (A) 7-week-old wild-type (Col-0) and *hipp* plants. (B) The primary inflorescence height of plants at the end of the life cycle. (C) Inflorescence stem (1 cm above the rosette level) from the 4-week-old plants. Scale bars = 1 mm. (D) Inflorescence stem diameter measured 1 cm above the rosette level. Data are means ± SD (*n* = 5). (E) Number of axillary rosette branches. Data are means ± SD (*n* = 15). (F) Representative pictures illustrating the increased rosette branching of *hipp32/33/34*. Different letters indicate individual groups for multiple comparisons with significant differences (one-way ANOVA, Wilcoxon, *p* < 0.05).

**Figure S4.** *hipp* mutants show abnormal flower development. (A) Inflorescences of the 5-week-old wild-type (Col-0) and *hipp* plants. (B) Silique formation rate (C) number of siliques on the main inflorescence stem at the end of a life cycle. (D) Time of flowering termination. (E-F) Phenotypes of flowers at different developmental stages. Staging according to (Smyth *et al*., 1990). Data are means ± SD (*n* = 15). Different letters indicate individual groups for multiple comparisons with significant differences (one-way ANOVA, Wilcoxon, *p* < 0.05)

**Figure S5**. HIPPs regulate root system architecture. Quantification of the (A) primary root length, (B) lateral root number, and (C) lateral root density in the 11-day-old wild-type (Col-0) and *hipp* plants. Data are means ± SE (*n* ≥ 10). Different letters indicate individual groups for multiple comparisons with significant differences (one-way ANOVA, Wilcoxon, *p* < 0.05).

**Figure S6**. *hipps* show increased sensitivity towards exogenous cytokinin. (A) Confocal images of the *TCSn:GFP* reporter gene (green) in roots of the wild-type (Col-0) and *hipp* plants treated with 1 µM BA or Mock (0.001% DMSO) for 16 h. Propidium iodide (PI, magenta) was used for cell visualization. Scale bars = 25 µm (B) Quantification of the *TCSn:GFP* fluorescence in white boxes in (A). Data are means ± SE (*n* ≥ 5). Different letters indicate individual groups for multiple comparisons with significant differences (one-way ANOVA, Wilcoxon, *p* < 0.05).

**Figure S7**. Cytokinin content is not changed in the *hipp* mutants. The total content of different cytokinin (CK) types in the 11-day-old wild-type (Col-0) and *hipp* seedlings. Data are means ± SD (*n* = 5). Asterisks indicate significant differences (one-way ANOVA; Dunnett test ; \**p* < 0.05, \*\**p* < 0.01, and \*\*\**p* < 0 .001).

**Figure S8**. Top 10 enriched GO terms in the (A) *hipp32*, (B) *hipp33*, and (C) *hipp34* plants.

**Figure S9.** Top 5 enriched KEGG pathways in the (A) *hipp32,* (B) *hipp33,* and (C) *hipp34* plants.

**Figure S10.** The HIPP34 HMA domain and the isoprenylation motif are relevant for PD localization. Subcellular localization of (A) GFP-HIPP34, (B) GFP-HIPP34^hma^, and (C) GFP-HIPP34^C470G^ co-expressed transiently in *N. benthamiana* leaves.

**Figure S11.** Spatiotemporal expression pattern of *pHIPP32*:*GUS*. *pHIPP32*:*GUS* reporter is expressed (A) in the radicle and colette regions of mature embryo, (B) in the shoot apex at the base of young leaves, at the root-hypocotyl junction, within the differentiated root vascular tissues and in the root apical meristem of a 5-day-old seedling. In 7-day-old seedlings, the *pHIPP32*:*GUS* activity is present (C) in the root meristem, (D) in the root vasculature, and in the emerging lateral root and (E) at the margins of the young leaves. In older plants, *pHIPP32:GUS* activity is observed (F) in the vasculature and stomata of rosette leaf and (G) in the stem vascular tissue. In flowers, strong *pHIPP32:GUS* expression is detected (H) in the flower stalks, (I) in the carpel walls of the gynoecium, and (J) in the vascular bundle of the anthers. In the fruit, GUS expression appears exclusively at (K) the silique septum. Scale bars = 100 μm; exceptions: B, I = 1 mm and H = 3 mm.

**Figure S12.** Spatiotemporal expression pattern of *pHIPP33:GUS*. In the 5-day-old seedlings, p*HIPP33*:GUS expression is localized (A) in the stipules of arising leaves and (B) in the quiescent center cells and in the columella stem cells of the root. In the 7-day-old seedlings, *pHIPP33:GUS* activity is detectable (C) in the stipules, in the shoot apical meristem (arrow) and (D) in the vasculature of the hypocotyl. (E-I) In flowers, *pHIPP33:GUS* expression is present in the gynoecium of different stages. Scale bars = 100 μm; exceptions: A and E = 1 mm.

**Figure S13.** Spatiotemporal expression pattern of *pHIPP34:GUS*. (A) Mature embryos expressing *pHIPP34:GUS* in the cotyledons, epidermis and in hypocotyl provasculature. Strong GUS expression is observed at (B) the shoot apex, in (C) tissues along the vasculature axis from hypocotyl to the root apical meristem and in (D) the vascular tissue of the emerging lateral root. *pHIPP34:GUS* activity is detectable in young leaves of (E) 5-day-old and (F) 7-day-old seedlings. In rosette leaves, *pHIPP34:GUS* is expressed in (G) the vasculature and in (H) the guard cells. (I) In flowers, *pHIPP34:GUS* is expressed in the pedicels, petals, in the style and the gynophore of the gynoecium, as well as in the filaments of the anthers. In siliques, *pHIPP34:GUS* activity is confined to (J) the outer seed integuments. Scale bars = 100 μm; exceptions: B, F, I = 1 mm and G = 5 mm.

**Table S1.** Cytokinin metabolite content in the 11-day-od wild-type (Col-0) and *hipp* plants expressed in pmol/ g fresh weight. iP, N6-(Δ2-isopentenyl)adenine; iPR, N6-(Δ2-isopentenyl)adenosine; iPRMP, N6-(Δ2-isopentenyl)adenosine 5’-monophosphate; iP7G, N6-(Δ2-isopentenyl)adenine 7-glucoside; iP9G, N6-(Δ2-isopentenyl)adenine 9-glucoside; tZ, trans-zeatin; tZR, trans-zeatin riboside; tZRMP, trans-zeatin riboside 5’-monophosphate; tZOG, trans-zeatin O-glucoside; tZROG, trans-zeatin riboside O-glucoside; tZ7G, trans-zeatin 7-glucoside; tZ9G, trans-zeatin 9-glucoside; cZ, cis-zeatin; cZR, cis-zeatin riboside; cZRMP, cis-zeatin riboside 5’-monophosphate; cZOG, cis-zeatin O-glucoside; cZROG, cis-zeatin riboside O-glucoside; cZ7G, cis-zeatin 7-glucoside; cZ9G, cis-zeatin 9-glucoside. Significant differences to Col-0 were evaluated by one-way ANOVA followed by the Dunnett post hoc test (*0.01 < *p* < 0.05, **0.001 < *p* < 0.01 and \*\*\**p* < 0.001. Values represent means ± SD (*n* = 5).

**Table S2**. List of primers used in this study.

